# Tough and Tunable Scaffold-Hydrogel Composite Biomaterial for Soft-to-Hard Musculoskeletal Tissue Interfaces

**DOI:** 10.1101/2020.01.22.915850

**Authors:** Raul A. Sun Han Chang, Mariana E. Kersh, Brendan A.C. Harley

## Abstract

Biological interfaces connecting tissues with dissimilar mechanical and structural properties are ubiquitous throughout the musculoskeletal system. Tendons attach to bone via a fibrocartilaginous interface (enthesis) that reduces mechanical strain and resultant tissue failure. Despite this toughening mechanism, tears at the enthesis occur due to acute (overload) or degradative (aging) processes. Repair involves surgical fixation of the torn tendon to bone, but results in the formation of a narrow fibrovascular scar tissue with inferior biomechanical properties. Progress toward enthesis regeneration requires biomaterial approaches to protect exogenously added or endogenously recruited cells from high levels of strain at the interface between dissimilar materials. Here, we describe an innovative reinforcement strategy to address this need. We report a stratified scaffold containing collagen bone and tendon tissue compartments linked by a continuous polyethylene glycol (PEG) hydrogel interface. Tuning the gelation kinetics of the hydrogel modulates its integration with the surrounding biomaterial compartments and yields biomechanical performance advantages. Notably, the continuous hydrogel interface reduces the deleterious effects of strain concentrations that form between tissue compartments in conventional stratified biomaterials. This design of mechanically robust stratified composite biomaterials may be appropriate for a broad range of tendon and ligament-to-bone insertions.

---

The enthesis is a stratified fibrocartilaginous tissue (250-500μm wide) that contains gradients in cell phenotype, biochemical cues, mineral content, as well as matrix composition and alignment.^[1]^ Importantly, this unique interfacial tissue microenvironment facilitates functional load-bearing by providing a continuous energy-absorbing zone of high compliance, an important tissue toughening mechanism under tensile loads.^[2]^ The classic enthesis injury is the rotator cuff tear, where acute overload, degeneration with age, or a combination of the two leads to partial or full-width tears within the tendon-to-bone enthesis. Surgical reattachment of tendon to bone is the clinical standard, but leads to formation of narrow fibrovascular scar tissue rather than a graded fibrocartilage enthesis. The resultant sharp boundary between mechanically mismatched tendon and bone leads to strain concentrations that significantly increases the risk of re-failure (>90% in some older demographics).^[3]^ Functional reintegration of the torn tissues requires regeneration of the compliant fibrocartilaginous interface, however, progress towards regenerative strategies for enthesis repair is hampered by a lack of biomaterial designs able to meet the unique functional requirements of these tissues.

Stratified biomaterials offer potential advantages for enthesis repair. Such biomaterials may selectively present optimized patterns of signals with features such as composition, structure, and mechanics tailored within discrete regions to spatially-regulate cell bioactivity and tissue remodeling. This stratified, spatially-controlled biological response is essential to recapitulate the distinct tissue microenvironments across the enthesis. To this end, we recently described a lyophilization method to fabricate biphasic collagen scaffolds containing tendon (anisotropic) versus bone (mineralized) compartments with distinct composition and microstructure connected by a continuous interface.^[4]^ The mineralized osseous compartment promotes mesenchymal stem cell (MSC) osteogenic differentiation and improves bone regeneration without supplemental osteogenic factors. Similarly, a non-mineralized anisotropic tendinous compartment promotes transcriptomic stability of primary tenocytes and induces MSC tenogenic differentiation.^[4, 5]^ To date, most biomaterials for tendon-to-bone enthesis repair, including this biphasic scaffold, replicate the enthesis as a gradient transition between tendon and bone rather than a unique multi-scale tissue. While significant progress has been made in biomaterials for tendon and bone repair, the inherent mismatch between tissue analogous compartments can be biologically and mechanically detrimental.^[6]^ Mechanical loading is required for the development and maintenance of tendon, bone, and the enthesis, and is unavoidable following injury and repair.^[1, 7]^ In tendon-bone biomaterials, mechanical mismatch at an interface between inherently dissimilar materials leads to strain concentrations under physiological loading that can significantly reduce cell viability and become a likely point of fracture. As such, the resulting mechanical strain that occurs between tendinous and osseous material compartments leads to a dampened cellular response and graft failure at the precise location where regeneration is needed.

Here we report development and validation of a unique biomaterial reinforcement motif inspired by structure-function properties of the native enthesis: inclusion of a compliant hydrogel interface between mechanically mismatched tendinous and osseous biomaterials.^[2, 8]^ While common in engineering materials, such design elements have not been previously explored in tissue engineering biomaterials. Here, we have developed an approach to control the insertion and stabilization of a compliant hydrogel zone between the tendinous and osseous scaffold compartments of our previously described biphasic collagen biomaterial. The resulting triphasic biomaterial is different from layered 2-phase or 3-phase biomaterials that lack a continuous interface. We show that tuning the fabrication parameters of this hydrogel zone provides a robust method to reduce levels of strain concentrations that form between dissimilar tissue compartments. Inclusion of a hydrogel enthesis also dramatically improves the macro-scale mechanical performance of the entire biomaterial and provides a new paradigm for tissue engineering approaches to improve healing for a wide range of musculoskeletal tissue insertions.

Stratified scaffolds can be created by fusing individually processed materials into a single construct or by simultaneously processing and integrating different phases during fabrication.^[9, 10]^ We previously developed a process to create biphasic scaffolds via liquid layering, where diffusive mixing between mineralized and non-mineralized collagen suspensions followed by lyophilization creates a porous scaffold with distinct phases but a continuity of collagen fibers across a distinct interface. While this interface resists delamination to an extent, under tension the interface is subject to high strain and eventual fracture, as is the case in similar scaffolds incorporating mechanically dissimilar phases.^[10]^ Here we hypothesize a similar layering and fusing approach could be employed with a third, enthesis-specific interfacial polyethylene glycol (PEG) hydrogel. PEG has been employed for a wide range of tissue engineering applications due to its non-cytotoxicity and the diverse functional groups that can be added to its backbone to facilitate crosslinking, degradability, improved bioactivity, and biomolecular functionalization, thereby providing a canvas for locally presenting cues to alter cellular response.^[11]^ We further hypothesize the rate of gelation of the PEG hydrogel can be used to control the extent of diffusive incorporation of the three different phases into a continuous scaffold.

To optimize design principles resulting in stable triphasic scaffolds, we optimized the gelation kinetics of the interfacial hydrogel phase. We employed a horseradish peroxidase (HRP)-mediated chemical polymerization to covalently crosslink 4-arm PEG-thiol (PEG-SH) monomers (**Figure 1A**).^[12]^ This class of reaction introduces tunability in crosslinking rate and hydrogel material properties (e.g. elasticity), both of which can be quantified via small amplitude oscillatory shear (SAOS) rheometry through conventional (*t*_*cross*_ and *G’*_*eq*_) and novel (*Δt*_*gel*_) parameters of gelation (**Figure S1**).^[13, 14]^ Here, *t*_*cross*_ defines the time it takes to transition from a predominantly viscous (*G’*<*G’’*) to an elastic (*G’*>*G’’*) material, while the equilibrium storage modulus (*G’*_*eq*_) is a measure of material elastic response. Recently, we reported the use of *Δt*_*gel*_, derived from the time derivative of *G’*, to quantify the duration over which significant changes in viscoelastic properties occur, from the onset of measurable gelation to reaching an equilibrium gel state.^[14]^ We selected a test set of PEG-SH hydrogels spanning gelation properties to match the time-scale for lyophilization based scaffold fabrication (*t*: 0-60min)^[15]^ with a range of elastic behavior (*G’*_*eq*_: 4-15kPa) from a library of previously characterized hydrogels (**Table S1**).^[14]^

**Figure 1.**
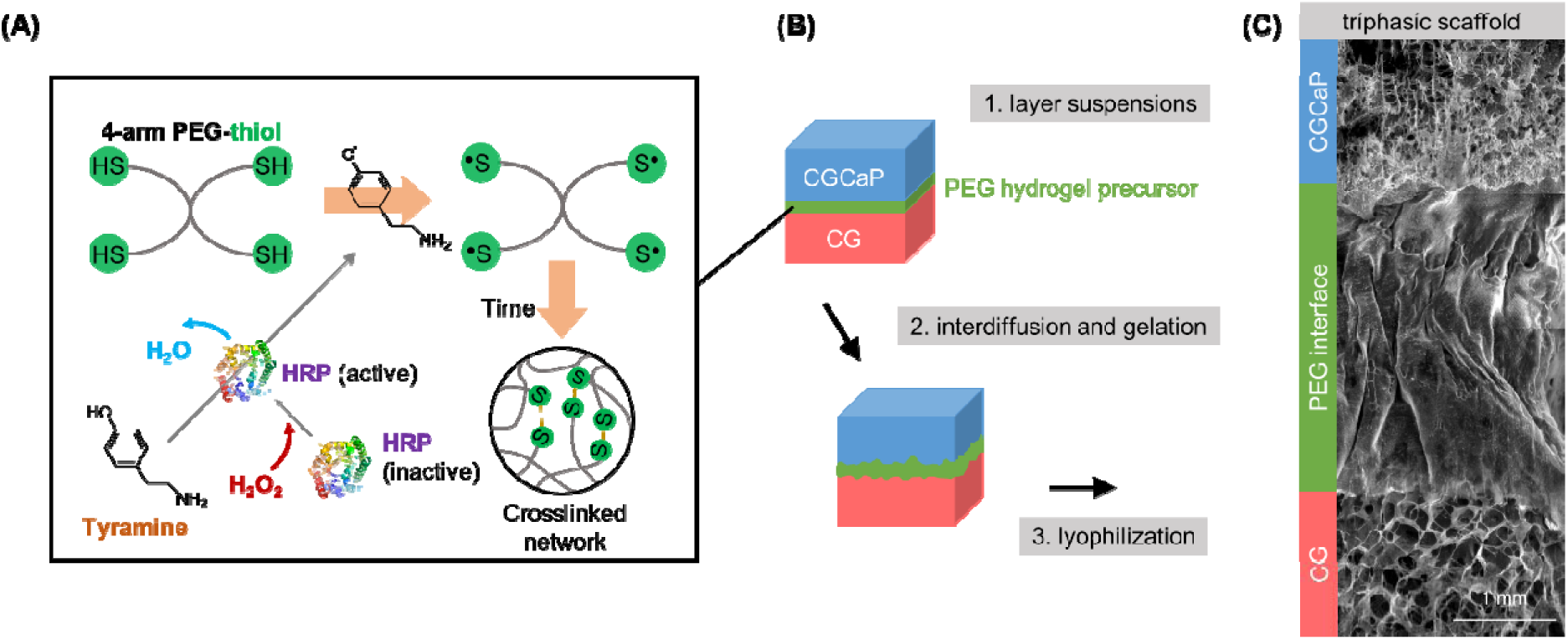
**A)** Formation of a crosslinked PEG network via horse radish peroxidase (HRP) catalyzed cross-linking. Initially, hydrogen peroxide (H_2_O_2_) reacts with HRP in its inactive state. Activated HRP oxidizes tyramine to form phenolic radicals that will oxidize thiol groups. Thiol groups on 4-arm PEG-thiol (PEG-SH) monomers are oxidized to thiol radicals that readily form disulfides over time to create a crosslinked polymer network. **B)** A suspension-layering lyophilization method is used to incorporate a PEG hydrogel layer between tendinous (CG) and osseous (CGCaP) collagen-GAG compartments. First, CG and CGCaP liquid suspensions and the PEG hydrogel precursor solution are layered into a mold and given time for diffusive mixing at their interface as the PEG precursor solution gels. **C)** Following lyophilization, structurally continuous triphasic scaffolds are generated with a distinct interfacial PEG hydrogel layer between GG and CGCaP compartments.

To fabricate the final triphasic scaffolds, we adapted a copper-polytetrafluoroethylene (PFTE) mold (**Figure S2**) that allows horizontal loading of liquid suspensions for precise scaffold layers and controlled phase interdiffusion prior to lyophilization (**Figure 1B**). After lyophilization, structurally continuous triphasic scaffolds are formed with a PEG hydrogel interfacial layer connecting the tendinous and osseous scaffold compartments (**Figure 1C**). The thermal conductivity mismatch between copper and PTFE at one end of the mold establishes a localized directional solidification environment to induce formation of an anisotropic, non-mineralized (tendinous) scaffold compartment, while at the opposing end an isotropic mineralized (osseous) collagen scaffold structure is formed (**Figure S3**).

ESEM images of resultant triphasic scaffolds demonstrate the topology and extent of incorporation of the interfacial hydrogel seam can be adapted via hydrogel gelation parameters. Overall faster gelling hydrogels (fast *t*_*cross*_ and short *Δt*_*gel*_) are more uniformly incorporated into scaffolds in distinct, monolithic hydrogel layers, whereas the slowest gelling hydrogel is distributed within the collagen fibers (**Figure 2A**), likely due to extended diffusive mixing between the hydrogel and collagen suspensions. The width of the scaffold hydrogel interface is also controlled via gelation (**Figure 2B**), giving us the ability to fabricate triphasic scaffolds with distinct physical characteristics based on a range of incorporated interfacial hydrogels. We report the mechanical performance of a series of triphasic scaffolds based on hydrogel gelation characteristics. Hydrogel interfaces were classified as having: fast (3-4 min), medium (6 min), or slow (12 min) *t*_*cross*_; short (7-9 min) or long (31-51 min) *Δt*_*gel*_; and low (1-5 kPa) or high (10-15 kPa) *G’*_*eq*_ (e.g., fast:long:high).

**Figure 2.**
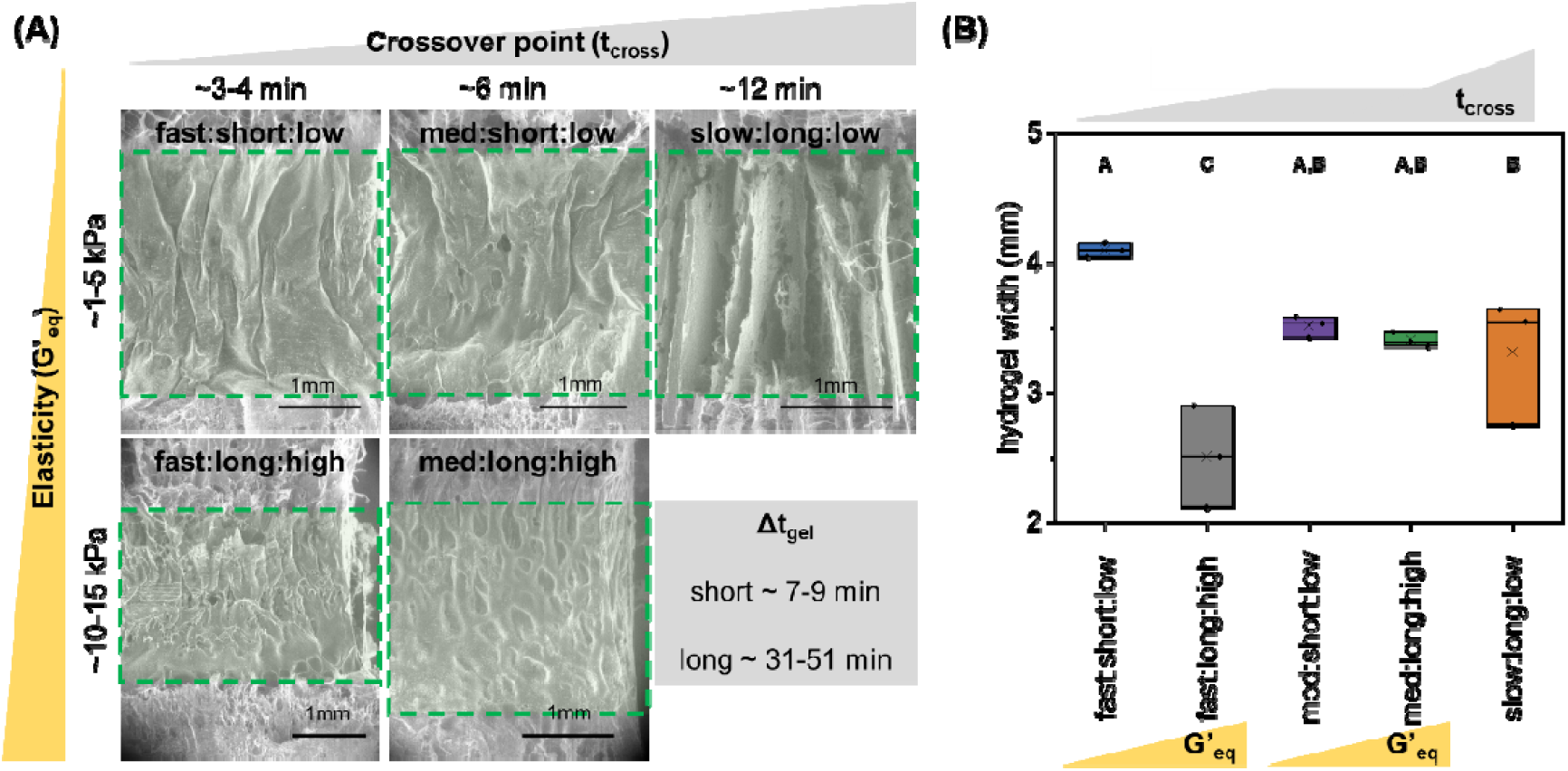
**A)** Representative ESEM images of triphasic scaffolds show that the extent of incorporation and topology of the interfacial hydrogel (green dashed-line region) is dependent upon the gelation characteristics (*t*_*cross*_, *G’*_*eq*_, and *Δt*_*gel*_) of the hydrogel. **B)** The width of the hydrogel layer in triphasic scaffolds is a function of its gelation characteristics (groups not sharing a letter are significantly different (p□<□0.05)).

We demonstrated gelation-dependent incorporation of a compliant hydrogel seam is an effective toughening mechanism in scaffolds under tension (**Figure S4**). Bulk scaffold toughness (area under the stress-strain curve) was significantly increased in triphasic scaffolds containing fast gelling (*t*_*cross*_) hydrogel formulations (fast:long:high and fast:slow:low) compared to biphasic scaffolds that lack a hydrogel insertion or triphasic scaffolds with slower *t*_*cross*_ (**Figure 3A**). Toughness appeared not to be strongly influenced by overall gelation time (*Δt*_*gel*_) or elastic properties (*G’*_*eq*_) of the hydrogel phase. Fast gelling enthesis variants display significantly higher toughness than other triphasic variants (e.g. med:long:high), indicating that incorporation of a compliant hydrogel interface alone is not sufficient to improve toughness.

**Figure 3.**
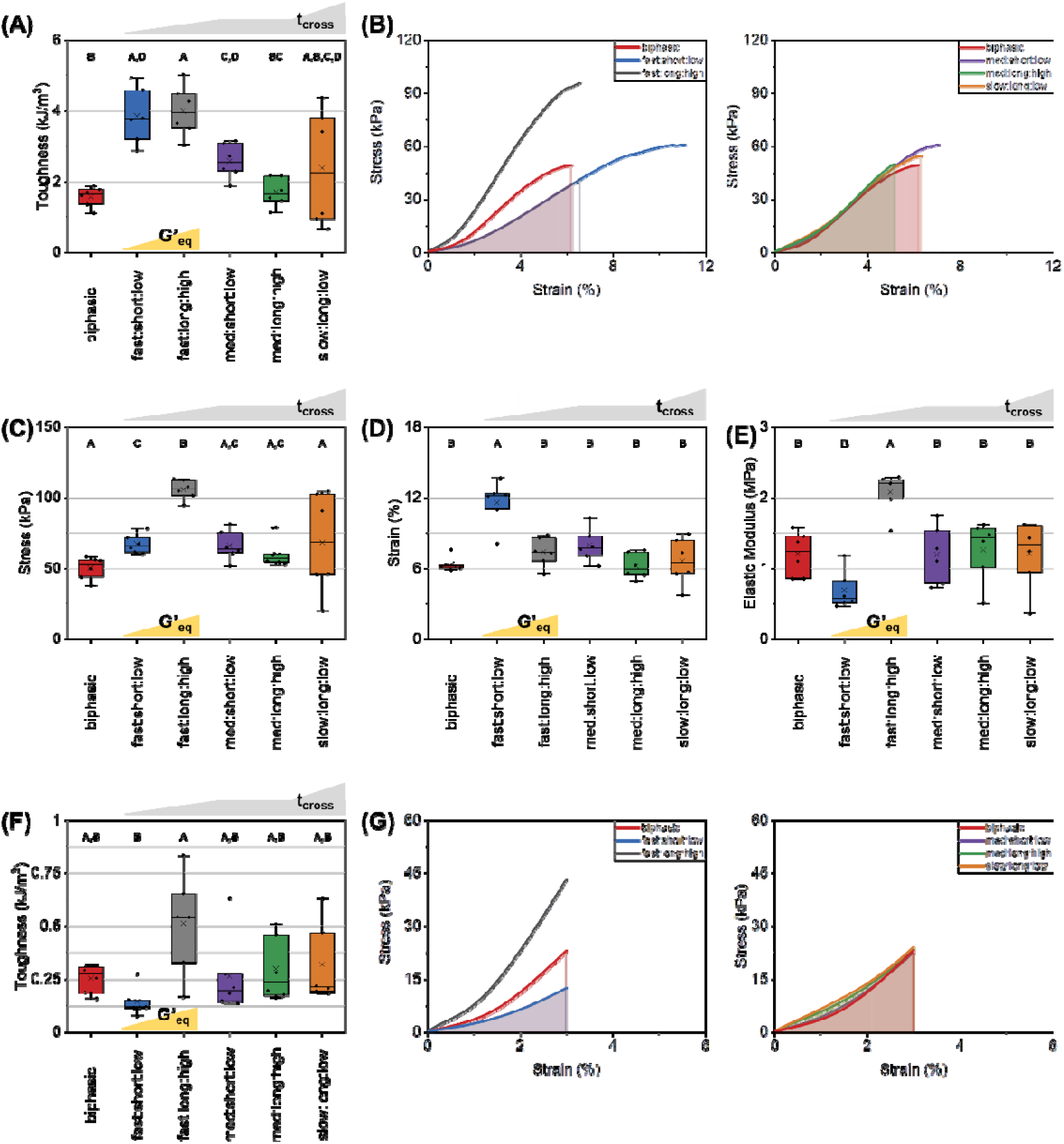
**A)** Bulk scaffold toughness through the point of failure. **B)** Averaged stress-strain curves for highest-toughness triphasic scaffold variants (left) and all other triphasic variants (right) versus the biphasic scaffold. **C)** Maximum tensile stress and **D)** strain at scaffold fracture. **E)** Bulk elastic modulus of scaffolds. **F)** Bulk scaffold toughness up to physiological levels of strain (3%). **G)** Averaged stress-strain curves for highest-toughness triphasic scaffold variants (left) and all other triphasic variants (right) versus the biphasic scaffold up to physiological levels of strain (3%). For each figure, groups not sharing a letter are significantly different (p□<□0.05). For stress-strain curves, linear interpolation was used to average multiple stress-strain curves for each scaffold.

Tuning gelation parameters of the incorporated hydrogel phase directly modulates resultant mechanical properties. Hydrogels that quickly become viscous (3-4 min), regardless of time required to fully gel (*Δt*_*gel*_), optimally integrate between flanking collagen suspensions during liquid phase layering and form mechanically robust triphasic scaffolds. Furthermore, differences in hydrogel storage modulus between the highest-toughness variants (fast:long:high and fast:short:low) did not directly influence fracture toughness, indicating that within the range of tested *G’*_*eq*_ (5-15 kPa), matching the hydrogel gelation with diffusive incorporation and lyophilization timescales is the key determinant of scaffold toughness. However, the elastic properties of the hydrogel phase may control how toughening of the scaffold occurs. Interestingly, the mechanism by which highest-toughness scaffolds increase toughness differs (**Figure 3B**). Some interfaces (fast:long:high) significantly increase maximum stress withstood (**Figure 3C**) and elastic modulus (**Figure 3E**), resulting in a steeper stress-strain curve and increased maximal stress. Comparatively, fast:short:low variants significantly increase strain tolerated prior to fracture (**Figure 3D**), and display a significantly lower elastic modulus but higher ductility. These distinct toughening mechanisms become significant under applied physiological strains (max. 3% applied strain; **Figure 3F, 3G**), where fast:short:low variants that display increased ductility show significantly lower levels of toughness than higher stiffness fast:long:high enthesis variants.

The inclusion of a hydrogel enthesis significantly alters the mode of failure and the local strain experienced by the scaffolds under tensile loading. Grossly, biphasic scaffolds that lack a hydrogel enthesis fail at the tendinous-osseous interface, while triphasic variants that display increased toughness fracture in one of the collagen compartments and away from the scaffold interface. Comparatively, triphasic variants that did not exhibit increases in bulk toughness also fail at the interface. To further characterize local changes in strain distribution underlying these bulk responses, we used digital image correlation (DIC) to map local strain on full-length scaffolds under tension (**Figure 4A**).^[16, 17]^ At 3.3% bulk applied strain, biphasic scaffolds display concentrated strain (∼10%) at the interface between the two disparate compartments, which ultimately fractures at 3.5% bulk applied strain. Triphasic variants that do not display improved toughness show similar strain concentrations. For example, the med:long:high variant displays ∼5% strain at the interface in response to 2.2% bulk applied strain, and ultimately fractures at 2.5% bulk applied strain. However, the highest-toughness triphasic variants display significantly reduced strains at the interface with heightened strain in the more elastic tendinous compartment, where it ultimately fractures at 8.7% and 7.4% bulk applied strain. We subsequently examined the strain profiles within the transition zones between tendon and bone scaffold compartments (**Figure 4B**). Again at 3.3% bulk applied strain, the biphasic scaffold interface develops sharp strain concentrations (7.2%), whereas highest-toughness triphasic scaffolds show evenly distributed interfacial strains (∼2%) that are less than the overall bulk applied strain. Clearly, tuning hydrogel gelation parameters to optimize integration of a compliant interfacial zone provides a facile way to improve overall mechanical performance and reduce localized strain concentrations between mechanically dissimilar biomaterial compartments.

**Figure 4.**
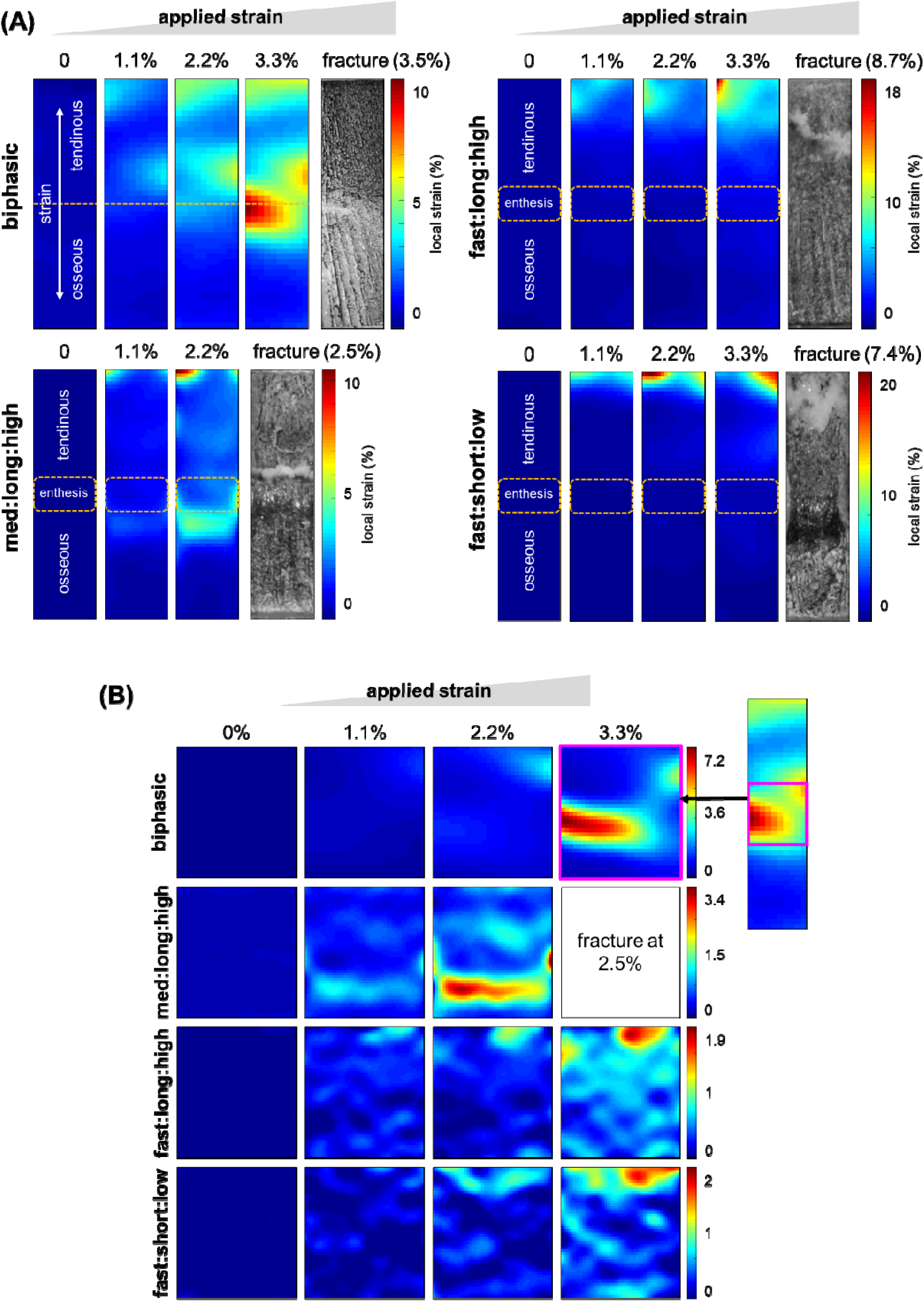
**A)** Representative profiles of local strain across the entire scaffold and **B)** within the middle regions of biphasic, med:long:high, fast:long:high, and fast:slow:low scaffolds at global applied strains of 0%, 1.1%, 2.2% and 3.3% (strains are with respect to the full-length scaffolds).

Ultimately, we demonstrate fabrication and validation of a novel class of tough, stratified biomaterials for repair of enthesis injuries. Inclusion of a compliant hydrogel interface between dissimilar collagen scaffolds provides a bioinspired reinforcement approach to effectively dissipate local strains between dissimilar biomaterial environments and reduce the prevalence of failure at the interface. We show that altering hydrogel gelation properties, notably the time required for viscous to elastic transition and the final elastic properties of the hydrogel network, provides a powerful tool to improve the mechanical performance of the resulting triphasic biomaterial. This new model for tough composite biomaterials may offer insight regarding bioinspired toughening of stratified composite materials and design of robust tissue scaffolds for a range of orthopedic insertion injuries.

## Experimental Section

Detailed methods are outlined in the Supporting Information.

### Preparation of hydrogels and rheological analysis of gelation

The subset of hydrogels incorporated into triphasic scaffolds was selected from a previously created library of hydrogels characterized via small amplitude oscillatory shear (SAOS) rheometry.^[14]^

### Fabrication of scaffolds

Non-mineralized (CG) and mineralized (CGCaP) collagen-GAG suspensions and biphasic scaffolds incorporating both phases were prepared as previously described.^[4]^ Triphasic scaffolds incorporating CG and CGCaP phases with an interfacial PEG-SH hydrogel seam were fabricated using a copper-PFTE mold (**Figure S2**) enabling precise layering of the three phases and unidirectional heat transfer through the copper plate base during lyophilization.

### Scaffolds uniaxial tensile testing

Lyophilized scaffolds were embedded into polymer end-blocks such that the scaffold interface was at the center of the exposed gauge length. Test-ready embedded rectangular scaffolds were 5mm thick and 5mm wide with a 15mm gauge length. Uniaxial tensile testing was done using an Instron 5943 Mechanical Testing System with a 100 N electromechanical load cell (Instron, Norwood, MA). Scaffolds were gripped at their end-blocks to prevent slippage, a preload was set to remove any slack from scaffolds, and scaffolds were strained until failure. Elastic modulus was calculated as the slope of the linear elastic region of the stress-strain curve and toughness was calculated as the area under the stress-strain curve (**Figure S2B**).^[17, 18]^

### Mapping local strain across scaffolds using digital image correlation (DIC)

Scaffolds were prepared and embedded as described above. Embedded scaffolds were then speckle-patterned and underwent uniaxial tensile testing as described above. During testing, images were captured using a Canon EOS 5DS R DLSR camera with a Canon Macro 100mm (Canon, Tokyo, Japan). Sets of digital images taken during testing up to scaffold fracture were correlated using a version of the MATLAB file package “Digital Image Correlation and Tracking” (Copyright (c) 2010, C. Eberl, D.S. Gianola, S. Bundschuh) modified by Elizabeth Jones (Improved Digital Image Correlation version 4 – Copyright © 2013, 2014, 2015 by Elizabeth Jones) to calculate local strain across scaffolds.

## Supporting Information

Supporting Information is available from the Wiley Online Library or from the author.

## Acknowledgements

Research reported in this publication was supported by the National Institute of Diabetes and Digestive and Kidney Diseases of the National Institutes of Health under Award Number R01 DK099528 as well as the National Institute of Dental and Craniofacial Research of the National Institutes of Health under Award Number R21 DE026582. The content is solely the responsibility of the authors and does not necessarily represent the official views of the NIH. This work was supported by the Office of the Assistant Secretary of Defense for Health Affairs Broad Agency Announcement for Extramural Medical Research through the Award No. W81XWH-16-1-0566. Opinions, interpretations, conclusions and recommendations are those of the authors and are not necessarily endorsed by the Department of Defense. We are grateful for the funding for this study provided by the NSF Graduate Research Fellowship DGE-1144245 (RSHC). The authors are also grateful for additional funding provided by the Department of Chemical & Biomolecular Engineering and the Carl R. Woese Institute for Genomic Biology at the University of Illinois at Urbana-Champaign. A portion of this research was facilitated by equipment at the Imaging Technology Group within the Beckman Institute for Advanced Science and Technology at the University of Illinois at Urbana-Champaign.

Received: ((will be filled in by the editorial staff))

Revised: ((will be filled in by the editorial staff))

Published online: ((will be filled in by the editorial staff))

