## Supplemental Information for "Tough and Tunable Scaffold-Hydrogel Composite Biomaterial for Soft-to-Hard Musculoskeletal Tissue Interfaces"

Supplemental Methods

*Preparation of hydrogels and rheological analysis of gelation parameters:* The subset of hydrogels incorporated into triphasic scaffolds was selected from a previously characterized library of hydrogels.^[1]^ 4-arm PEG-thiol (PEG-SH) hydrogel precursor suspensions were generated from mixtures of 4-arm PEG-thiol (PEG-SH; 20,000 MW, JenKem Technology, Plano, TX, U.S.A) dissolved in DPBS (pH 7.3, Corning cellgro, Corning, New York, U.S.A) mixed with a reaction solution generated from H_2_O_2_ (30% solution, Macron, Radnor, PA, U.S.A), tyramine (99%, Sigma-Aldrich, St. Louis, MO, U.S.A), and horse radish peroxidase (HRP) (Thermo Fisher Scientific, Waltham, MA, U.S.A). All samples contained a fixed amount of tyramine (5mM) and HRP (5 units/mL). The polymer and reaction solutions were mixed to generate the polymerizing hydrogel solution. After mixing, samples were immediately loaded onto the bottom plate of a DHR-3 rheometer (TA Instruments, New Castle, DE, U.S.A) with a Peltier system controlling the temperature at 25 ± 0.1 °C. A parallel plate fixture (20 mm diameter) was used with a measuring gap of 1000 µm. Oscillatory shear was continuously applied at 0.8 rad/s at a sampling interval of 30 seconds. Mixing and loading times were added to the start of the measurements to account for the short period of reaction prior to starting the measurement.

*Preparation of collagen-glycosaminoglycan liquid suspensions:* Non-mineralized (CG) and mineralized (CGCaP) collagen-glycosaminoglycan suspensions were prepared as previously described.^[2]^ Briefly, CG suspensions consisted of 1 w/v% Type 1 collagen from bovine Achilles tendon (MilliporeSigma, Burlington, MA) and chondroitin-6-sulfate from shark cartilage (MilliporeSigma, Burlington, MA) at a ratio of 11.25:1, respectively, in 0.05M acetic acid. CGCaP suspensions incorporated calcium salts (Ca(OH)2, Ca(NO3)2∙4H2O) with the collagen and chondroitin-6-sulfate in a 0.1456M phosphoric acid/0.037M calcium hydroxide buffer solution to yield scaffolds with a calcium phosphate mineral content of 40 wt%.^[3]^ Both suspensions were homogenized at 7,000 rpm and 4°C to prevent collagen denaturation. Following homogenization, both suspensions were stored overnight at 4°C and degassed under vacuum prior to use.

*Fabrication of biphasic and triphasic scaffolds:* Biphasic scaffolds incorporating CG and CGCaP phases were fabricated as previously described.^[2]^ A mold comprising a polytetrafluoroethylene (PFTE) body joined to a 1/16” thick copper base was used to induce directional solidification due to the mismatch in thermal conductivity during the freeze-drying process. First, CG suspension (550µL) was pipetted into rectangular wells within the PFTE body (6 x 6mm area; 30 mm deep). CGCaP suspension (550µL) was then layered on top of the CG suspension, filling the well. The suspensions were then allowed to interdiffuse at room temperature for 20 min prior to freeze-drying. Triphasic scaffolds incorporating CG and CGCaP phases with an interfacial PEG-SH hydrogel seam were fabricated using a mold comprising 3 PFTE pieces, a 1/16” thick copper base plate, and a 3D-printed acetyl-butyl-styrene (ABS) phase divider (**Figure S2A**). In the loading conformation (horizontal) with a phase divider inserted, the mold cavity was filled with both degassed collagen-glycosaminoglycan suspensions (500µL each) and the polymerizing hydrogel solution (125µL) using a micropipette (**Figure S2B**). After the phase divider was removed, the 3 phases were allowed to mix diffusively at their interface for 20 minutes at room temperature. The mold was then switched to its lyophilization (vertical) conformation, whereby the significant mismatch in thermal conductivity between the PFTE body and copper base plate promotes unidirectional heat transfer through the copper base during lyophilization (**Figure S2C**). The suspension-loaded mold was placed into a freeze-dryer (VirTis, Gardiner, NY) held at 20°C for 5 min, then the temperature was ramped down to -10°C at a rate of 1 °C/min and held at -10°C for 2 h to complete freezing. Immediately following freezing, the shelf temperature was ramped up to 0 °C at a rate of 1 °C/min and scaffolds were dried via sublimation at 200mTorr. For measuring freezing kinetics during lyophilization, ABS shelves of 3 different lengths (7.5mm, 15mm, 22.5mm) were 3D-printed. Immediately after loading the mold cavity, one shelf is inserted into the cavity, and all subsequent steps carried out as before. Prior to lyophilization, a thermal probe is inserted into the shelf, with each shelf length corresponding to a scaffold phase based on its vertical location in the mold well (**Figure S3A**).

*Triphasic scaffold imaging and measurement of interfacial width:* Scaffolds were imaged using a FEI Quanta FEG 450 ESEM (FEI Company, Hillsboro, OR) at low vacuum. Interfacial hydrogel width was measured using the open source image processing package Fiji (National Institutes of Health, Bethesda, MD).^[4]^ Briefly, for each imaged triphasic scaffold, a reference scale was set (pixels/mm) based on the ESEM image scale. Then, the longest and shortest width of the hydrogel interface was measured, and the average of the two was reported as the hydrogel width for that scaffold.

*Preparation of scaffolds for uniaxial tensile testing:* Freeze-dried scaffolds were cut into a uniform length using a 3D-printed ABS cutting guide (21mm total length: 12.5mm from center to CG end, 8.5mm from center to CGCaP end). Both scaffold ends were then embedded into hollow 3D-printed ABS end-blocks such that the scaffold interface was at the center of the exposed gauge length (15mm) with equal lengths of CG and CGCaP exposed. End-blocks were used to protect scaffolds from being crushed by the pneumatic grips during mechanical testing. For the soft CG end, a two-component silicone rubber compound (RTV615; Momentive Specialty Chemicals Inc., Columbus, OH) at a ratio of 4:1 (A component: B component) was poured into a hollow end-block (5mm deep) and allowed to partially cure at room temperature for 60 min before inserting the scaffold end. For the rigid CGCaP end, super glue (Loctite, Düsseldorf, Germany) was used to adhere the scaffold end into a shallow (1mm deep) end-block. Mechanical testing was performed at least 24 hours after embedding to allow adhesive curing and ensure scaffolds would not pull out of the end-blocks during mechanical testing. Test-ready embedded rectangular scaffolds were 5mm thick and 5mm wide with a 15mm gauge length.

*Uniaxial tensile testing:* Uniaxial tensile testing was done using an Instron 5943 Mechanical Testing System with a 100 N electromechanical load cell (Instron, Norwood, MA). Scaffolds were held in place during testing by pneumatic grips at their end-blocks to prevent slippage. A preload was set to remove any slack from scaffolds, whereby scaffolds were strained at a rate of 0.1mm/min until a load greater than 0.05N was registered. Subsequently, the test would begin, and the scaffolds were strained at a rate of 1 mm/min until scaffold fracture. Elastic modulus was calculated as the slope of the linear elastic region of the stress-strain curve and toughness was calculated as the area under the stress-strain curve (**Figure S4B**).^[5, 6]^

*Mapping local strain on scaffolds under tension using digital image correlation (DIC):* Scaffolds were prepared and embedded as previously described. Embedded scaffolds were then speckle patterned with waterproof India ink (Black Art Materials, Galesburg, Illinois) using a gravity feed airbrush with a nozzle size of 0.3mm (GotHobby, South El Monte, CA). Scaffolds underwent uniaxial tensile testing. During testing, images were captured using a Canon EOS 5DS R DLSR camera with a Canon Macro 100mm lens at a rate of 1 image every 5 seconds using a time lapse remote (Canon, Tokyo, Japan). Scaffold fracture mode was shown as the final image captured after fracture occurred. Sets of digital images taken during testing up to scaffold fracture were correlated using a version of the MATLAB file package “Digital Image Correlation and Tracking” (Copyright (c) 2010, C. Eberl, D.S. Gianola, S. Bundschuh) modified by Elizabeth Jones (Improved Digital Image Correlation version 4 – Copyright © 2013, 2014, 2015 by Elizabeth Jones) to calculate local strain across scaffolds. The DIC file package calculates local strain with respect to a region of interest (ROI). The ROI is specified in the first image of the set, and displacement calculated from that first reference image for each subsequent image within the ROI up to scaffold fracture. The ROI was set as the entire exposed scaffold (**Figure 4A**) or encompassing just the scaffold middle zone (**Figure 4B**). Strain in **Figure 4B** was calculated with respect to the total scaffold gauge length by equating the ratio of the length of the scaffold interface ROI to the total scaffold gauge length and the ratio of strain with respect to the scaffold interface ROI to strain with respect to the total scaffold gauge length. For each set of scaffold images, reduced images were correlated first to generate initial guesses for displacements in full images using individually optimized correlation settings (image reduction factor: 7, subset size: 21-61; threshold: 0.5; search zone: 3, grid step size: 5-10). Reduced correlations were iterated up to 6 times for image sets with high displacements. Subsequently, full images were correlated using the reduced image data (subset size: 21-61; threshold 0.5; search zone: 2; grid step size: 5-10). Finally, displacements were smoothed prior to calculating strains to reduce noise (Gaussian distribution of weights; kernel size: 11; number of smoothing passes: 3; maximum size of contiguous non-correlated points to smooth over: 15) and local strains were calculated using a cubic (16-node) algorithm. Contour plots and line scans of strain could then be visualized. Line scan data was exported onto Microsoft Excel and average strain plotted as a function of position for each sample type.

*Statistical analysis: n*=3 samples for each hydrogel formulation were examined for rheological analysis of gelation, *n*=3 samples for each scaffold condition were examined for measuring interfacial hydrogel width, and *n*=6 samples for each scaffold condition were examined for uniaxial tensile testing. Statistical analysis was performed in RStudio (RStudio, Inc., Boston, MA) using the programming language R (R Core Team, Vienna, Austria). Normality of the residuals for all samples was verified using a Shapiro–Wilk test and homogeneity of variances was verified using Levene's Test. Statistical analysis of data satisfying normality and homogeneity of variances was performed using one-way analysis of variance (ANOVA) followed by Tukey-honest significant difference post hoc test. Statistical analysis of data not satisfying normality was performed using Kruskal–Wallis H test followed by Dunn's Multiple Comparison post hoc test. Statistical analysis of data not satisfying homogeneity of variance was performed using Welch’s ANOVA followed by Games-Howell post-hoc test. Statistical analysis of data not satisfying normality or homogeneity of variance was performed using a robust regression. Significance for all statistical analyses was set at p < 0.05.

Supplemental Figures

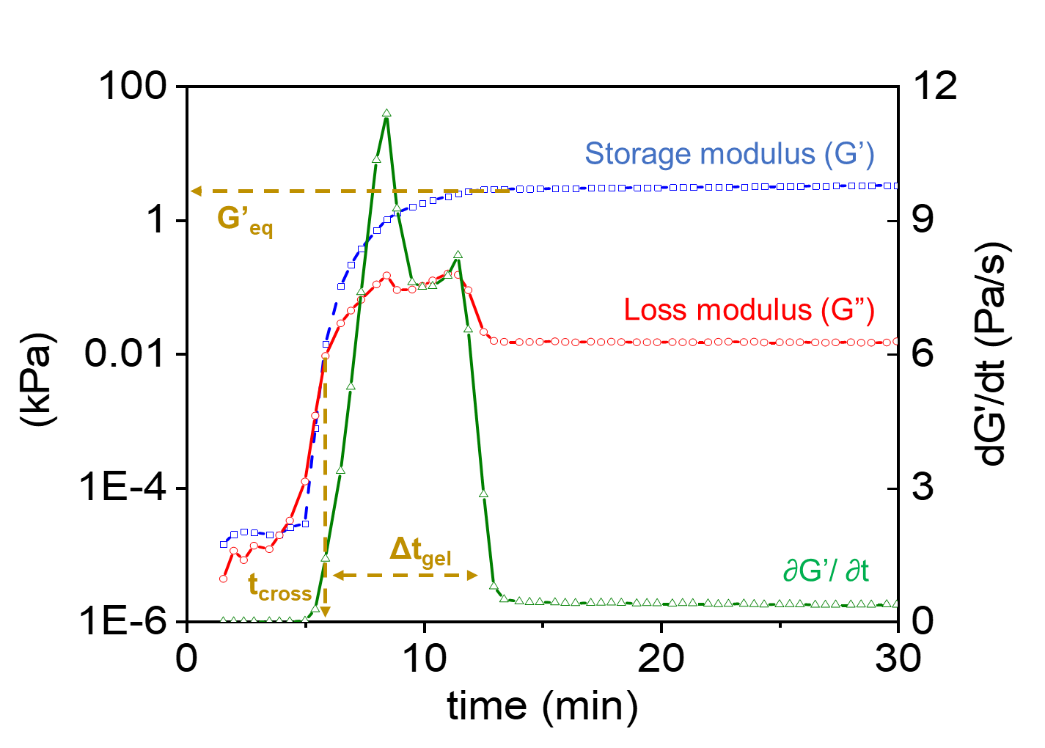

**Figure S1.** Small-amplitude oscillatory shear (SAOS) rheology at 0.8 rad/s and 2% strain was used to monitor the storage modulus (*G’*) and loss modulus (*G’’*) of crosslinking PEG-SH hydrogels as a function of time. For each hydrogel, the crossover state (*t_cross_*) was determined as the point where *G’* and *G’’* intersect (*G’*=*G’’*), the equilibrium storage modulus (*G’_eq_*) was obtained from the plateau value of *G’* at 60 minutes, and the gelation interval (*Δt_gel_*) was calculated as the full width at 10% of the maximum value of the derivative of *G’* (d*G’*/dT).

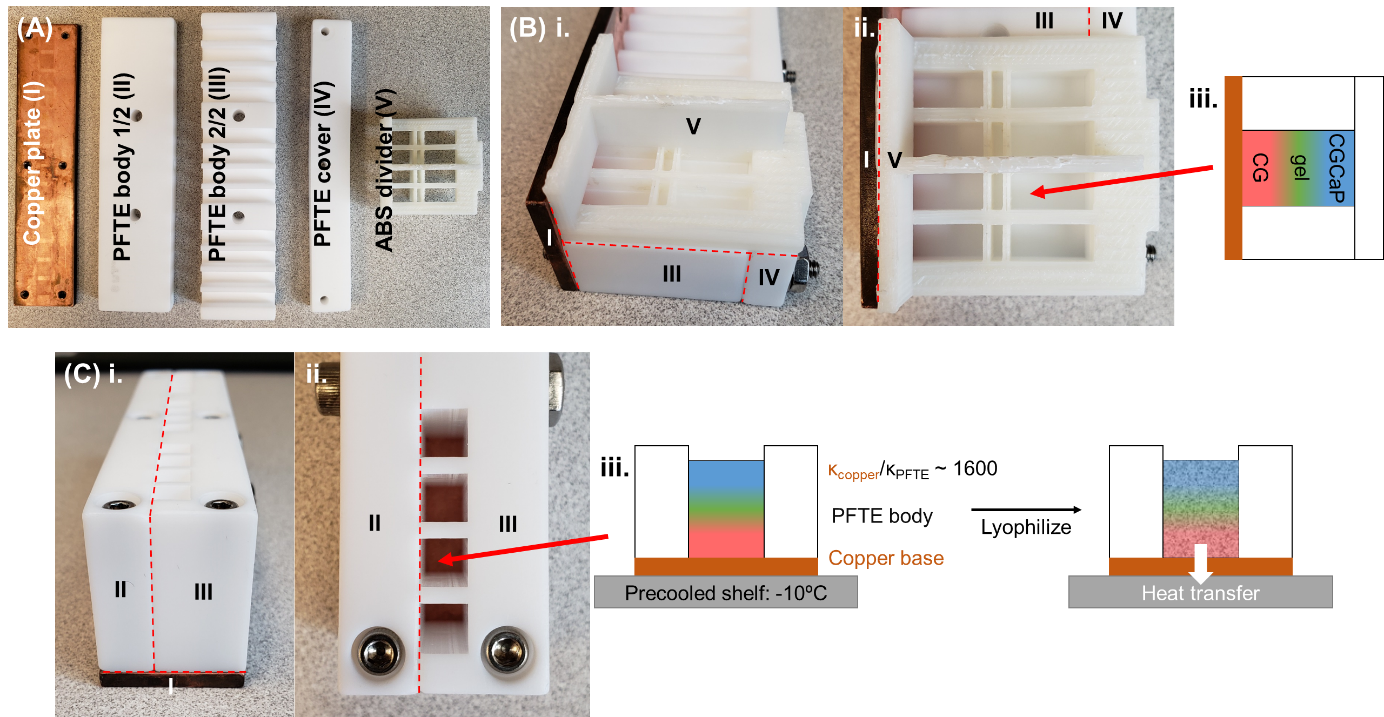

**Figure S2.** Schematic of mold used to fabricate triphasic scaffolds via lyophilization. A) The mold consists of 4 pieces and a divider. B) A phase divider is used to horizontally load liquid collagen and pre-gel suspensions into the mold cavity (i. side view and ii. Top view). iii. After the phase divider is removed vertically, the 3 liquid phases meet and mix diffusively at their interface until lyophilization. C) Following the phase mixing period (20 minutes at room temperature), the mold was sealed then placed into the freeze dryer in a vertical configuration: i. side view; ii. top view; iii. The significant mismatch in thermal conductivity between the polytetrafluoroethylene (PFTE) body and copper base plate promotes unidirectional heat transfer through the copper base during lyophilization.^[2]^

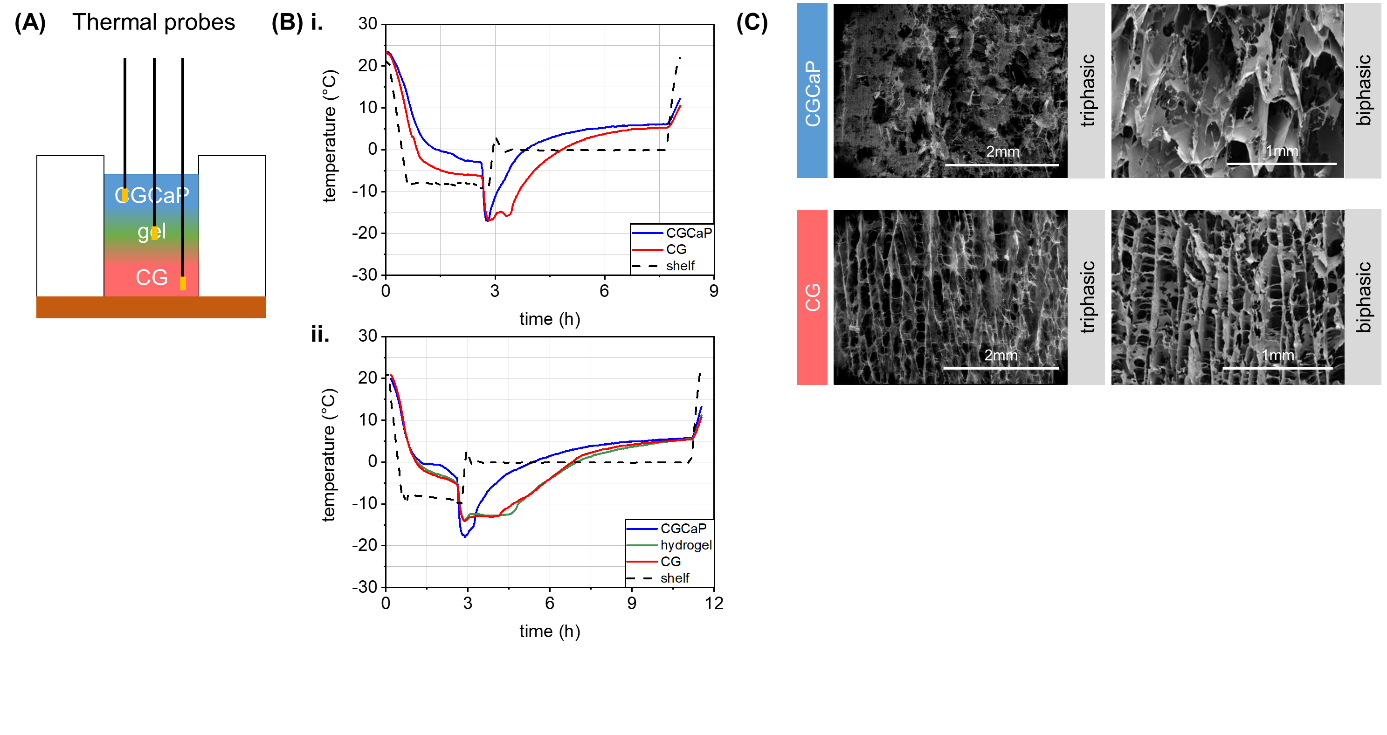

**Figure S3.** Monitoring local temperature profiles during lyophilization. A) Thermal probes were inserted at multiple depths using customized ABS support structures in the suspensions to monitor local freezing kinetics. B) Consistent temperature profiles were observed in the mineralized and non-mineralized collagen suspensions for both i. biphasic and ii. triphasic scaffolds (fast:short:low). C) Representative ESEM images of the resultant scaffold microstructure of both mineralized (CGCaP) and non-mineralized (CG) scaffold compartments in biphasic vs. triphasic scaffolds.

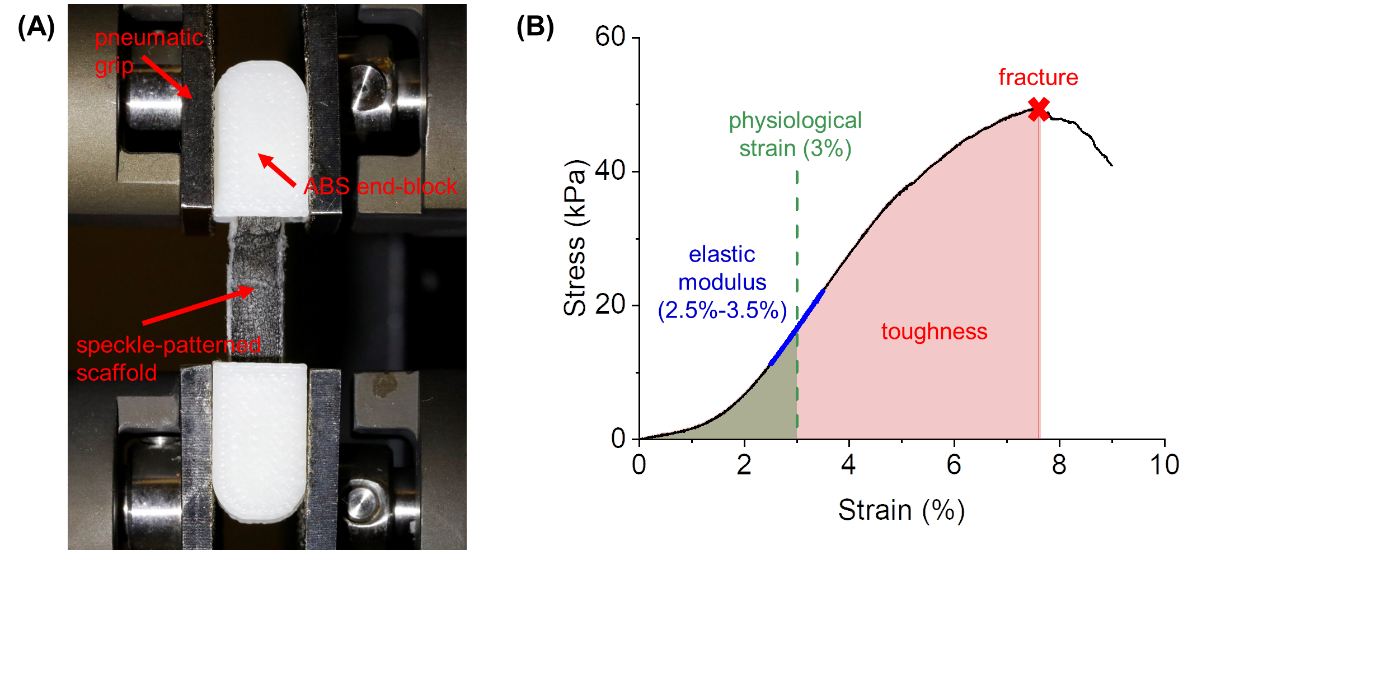

**Figure S4.** A) Uniaxial tensile testing was performed on speckle-patterned scaffolds embedded in ABS end-blocks. B) Scaffold mechanical performance was measured via tensile testing to generate stress-strain curves used to calculate elastic modulus (linear regime at 2.5%-3.5% strain^[6, 7]^) and toughness (area under stress-strain curve) up to scaffold fracture and physiological tendon strain (3%).^[8]^

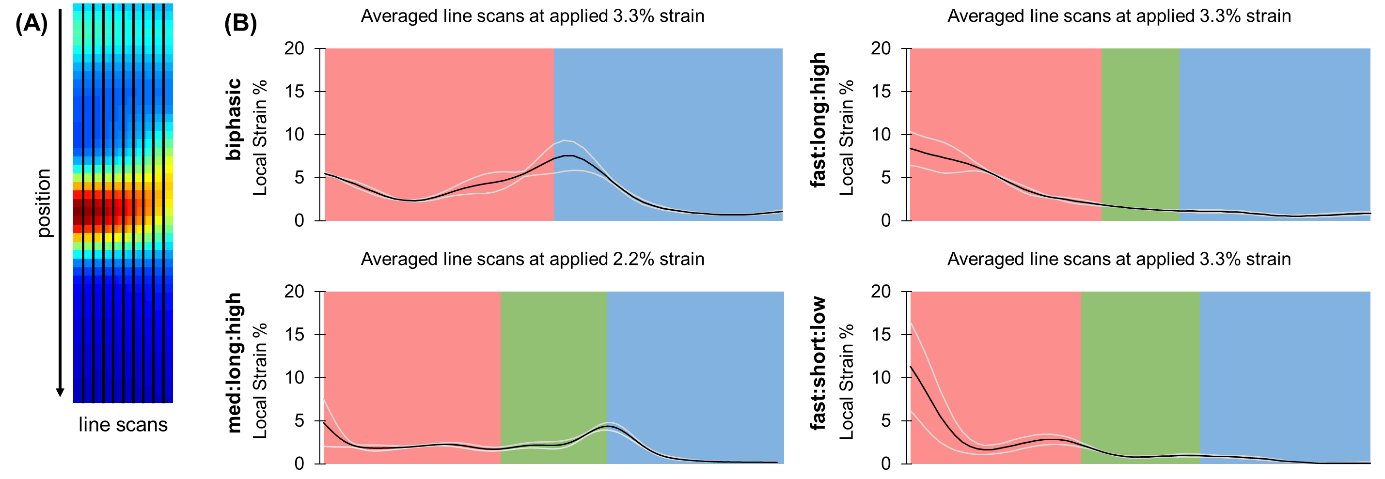

**Figure S5.** A) Mapping average strain profiles within scaffold variants as a function of position. Strain profiles were calculated using DIC data from 9 equally-spaced line scans along the length of the scaffold (parallel to direction of applied strain). B) The average (black line) and standard deviation (interval contained by gray lines) of strain magnitude along the entire length of each scaffold was reported at a global applied 3.3% strain (2.2% applied strain for med:long:high variant that fractured at 2.5% strain).

Supplemental Table

**Table S1.** A test set of hydrogel samples (varying PEG-SH (wt%) and H_2_O_2_ (mM) with tyramine (5mM) and HRP (5 units/mL) held constant) with a range of *t_cross_*, *Δt_gel_*, and *G’_eq_* was identified from previously reported SAOS measurements.^[1]^ The following naming convention was used to distinguish samples based on their gelling characteristics: time to reach crossover point (fast, med, slow), duration of gelation interval (short, long), and magnitude of *G’_eq_* (low, high). Reported values of *t_cross_*, *Δt_gel_*, and *G’_eq_* are average ± standard deviation (*n*=3).

| **Sample** | **PEG-SH (wt%)** | **H_2_O_2_ (mM)** | **t_cross_ (min)** | **Δt_gel_ (min)** | **G’_eq_ (kPa)** | **t_cross_: Δt_gel_: G’_eq_** |
| --- | --- | --- | --- | --- | --- | --- |
| P5H5 | 5 | 5 | 5.8±0.6 | 6.8±1.1 | 4.0±0.5 | med:short:low |
| P5H10 | 5 | 10 | 3.0±0.4 | 8.8±2.2 | 5.0±0.2 | fast:short:low |
| P10H5 | 10 | 5 | 11.9±2.7 | 51.2±12.9 | 1.0±0.4 | slow:long:low |
| P10H8 | 10 | 8 | 5.8±0.4 | 31.4±1.9 | 10.8±0.3 | med:long:high |
| P10H10 | 10 | 10 | 4.6±0.2 | 32.9±4.5 | 15.8±0.4 | fast:long:high |
